# Investigation of the Role of the LC3 Conjugation System in Autophagy for Human Reward System Reactivity

**DOI:** 10.1101/2025.03.26.645396

**Authors:** Jens Treutlein, Bernd Krämer, Oliver Gruber

## Abstract

**Objectives:** The dopaminergic reward system is involved in the etiology of psychiatric disorders, and autophagy has been suggested to interfere with dopamine release. LC3 conjugation plays a key role in autophagy and comprises modification of autophagy protein LC3 with phosphatidylethanolamine. We investigated whether LC3 conjugation may impact on the strength of activation in key regions of the mesolimbic reward system.

**Methods:** To test our hypothesis, responses of the reward system to conditioned stimuli were assessed using the ‘desire-reason dilemma’(DRD) paradigm, that allows for investigation of reward processing during fMRI. Association of a set of missense variants with reward system responses was analyzed in a sample of 214 participants.

**Results:** As a main finding, the gene set was associated with both ventral tegmental area (VTA) and nucleus accumbens (NAc) responses to conditioned reward stimuli (empirical P-value R-VTA: 0.008, R-NAc: 0.009). Strongest missense variants were *MAP1LC3B*_rs113610787 (P=3.219e-05) for association with response in the L-NAc, and *ATG4B*_rs143448469 (P=4.366e-05) for the R-NAc.

**Conclusions:** Findings provide evidence that variation of the LC3 conjugation system influences responses of the VTA and NAc to conditioned reward stimuli. Further studies are required to replicate the findings, and to investigate the possible role of LC3 conjugation in psychiatric disorders.

## Introduction

The LC3 conjugation system is long known to play a key role in the highly complex autophagy process and comprises the specific aspect of modification of autophagy protein LC3 with phosphatidylethanolamine (PE) (*Tanida et al. LC3 conjugation system in mammalian autophagy. Int J Biochem Cell Biol, 2004, 36, 2503-2518*). The underlying biochemical reactions constitute a phylogentically very old and conserved process, as lipidation of autophagy protein atg8, the ortholog of human LC3, occurs even in the yeast *Saccaromyces cerevisiae* (Thukral et al. 2015), an evolutionary extremely distant unicellular model organism.

In an initial biochemical reaction of the LC3-PE conjugation machinery, the cysteine protease ATG4B cleaves the carboxyterminal part of the amino acid chain of LC3 and generates a soluble form of LC3 (LC3-I) that is characterized by the amino acid glycine at the C-terminus, where then further modification of LC3-I occurs. The Gly120 residue at which the further molecular events for LC3 conjugation take place, is also conserved between humans and yeast (Thukral et al. 2015), which is the organism where mechanisms of autophagy were initially discovered and provided much of the current understanding of the biochemical processes (Ohsumi 2012, Ohsumi 2014).

In the further molecular events occurring in LC3 conjugation, soluble LC3-I is converted to lipid-bound LC3-II. To accomplish this, the autophagy protein ATG7 binds to LC3-I and recruits ATG3, which catalyzes a reaction that attaches (conjugates) the lipid phosphatidylethanolamine (PE) to LC3-I. This lipidation process is enhanced by binding of the ATG5-ATG12 complex that combines with ATG16L1 to form an oligomeric complex, which participates in the reaction. The conversion product of this reaction, lipidated LC3-II, is finally inserted into the membrane of the phagophore, a cup-shaped structure that engulfs cytosolic components during the autophagy process. Membrane-anchored LC3-II is essential for the elongation of the phagophore and as site for the attachment of autophagic cargo receptors (Besemer et al. 2021; Pfister 2023; Randall-Demllo et al. 2013; Yamamoto et al. 2023; Zhou 2022). The phagophore matures to an autophagosome which subsequently fuses with a lysosome to form the autolysosome, where hydrolases degrade the cellular material engulfed by the phagophore (Pfister 2023).

The overall process of autophagy balances the synthesis, recycling and degradation of cellular components, and thus plays a crucial role for the maintenance of protein homeostasis, also known as proteostasis, in neurons (Merenlender-Wagner et al. 2014; Yamamoto et al. 2023). Dysregulation of the selective degradation of synaptic components by autophagy causes proteostasis imbalances that have been shown to affect the function of the synapse at multiple levels, e.g., quality control of synaptic proteins (e.g., neurotransmitter receptors) and organelles, maintenance of synaptic plasticity, and regulation of synapse remodeling (Fleming and Rubinsztein 2020; Lieberman and Sulzer 2020; Limanaqi et al. 2018; Tomoda et al. 2020). Via these synaptic mechanisms, regulation of autophagy is thought to play a key role for the adaptation to stress (Peker and Gozuacik 2020; Limanaqi et al. 2020), contributes to the pathophysiology of psychiatric disorders (Merenlender-Wagner et al. 2015; Jia and Le 2015; Schneider et al. 2016; Tomoda et al. 2020; Tan et al. 2024), and impacts on the action of antipsychotics and antidepressants (Vucicevic et al. 2018; Rein 2019; Gan et al. 2024).

Among the neuronal populations of the CNS, autophagy was suggested to play a prominent role in dopamine neurons, as a modulator of dopaminergic neurotransmission. The interference of autophagy with the mechanism of dopamine release, and thus with dopamine neurotransmission, is thought attributable to increased damage of synaptic proteins by oxidation products of dopamine, and by altered trafficking of synaptic dopamine vesicles in these neurons (Munoz et al. 2012; Limanaqi et al. 2018). Populations of dopamine-containing neurons in the ventral tegmental area (VTA) that project to the nucleus accumbens (NAc) constitute the core of the mesolimbic dopamine pathway (Dichter et al. 2012; Russo and Nestler 2013). In humans, clear evidence for disturbances in the mesolimbic dopamine pathway for psychiatric disorders were provided by studies of reward-specific paradigms in functional magnetic resonance imaging (fMRI) studies (Deserno et al. 2016; Goya-Maldonado et al. 2016; Qi et al. 2021; Richter et al. 2015; Trost et al. 2014).

Testing the LC3 conjugation machinery for the aggregated contribution of genetic variants in groups of genes whose products play a role in this process is an approach to circumvent the extreme correction for multiple testing in single-marker analysis, under the assumption that functionally related genes contain multiple variants that disrupt function. An example for this are gene set-based methods (Wang et al. 2010), typically in the form of consecutive enzymes in a biochemical reaction or in the form of functional groups, by grouping genes whose protein products interact. Such a group of genes can be profoundly investigated using prioritization of functional sequence variants. Non-synonymous (missense) variants that change an amino acid in a protein to one with different physicochemical properties can have a very high risk for causing a phenotype, are known to account for many known disease associations, and therefore should be given high priority in candidate-gene association studies (Tabor et al. 2002).

We hypothesized that individuals carrying missense variants in genes whose products influence the LC3 conjugation machinery, display changes in subcortical reward processing in humans. For this purpose, we investigated effects of these genetic variants a sample of n=214 healthy young adults, using fMRI and genetic methods. We conducted gene set-based tests to investigate if the joint activity of these variants of the LC3 conjugation machinery influences the activation strength of brain regions within the reward system in humans.

## Materials and Methods

### Subjects

A homogeneous sample of n=214 participants (mean age ± SD: 24.12 years ± 2.465 years, minimum age: 19 years, maximum age: 31 years; 128 females and 86 males) underwent fMRI and was genotyped. The sample comprised adult young healthy individuals, as previously described (Trost et al. 2016). The ancestry of the participants was European.

The study was carried out in accordance with the Declaration of Helsinki and was approved by the local ethics committees, of the Medical Faculty of Göttingen University (number 14/3/09, date 02.07.2009) and of the Medical Faculty of Heidelberg University (number S-123/2016, date 09.03.2016). All participants provided written informed consent.

### Desire-Reason-Dilemma Paradigm

All subjects had performed the ‘desire-reason dilemma’ (DRD) paradigm, that allows for systematic investigation of systems-level mechanisms of reward processing in humans. In this fMRI experiment, participants initially undergo an operant conditioning task. Following this task, the DRD paradigm is performed by the subjects. For the present study, of the DRD paradigm one part was used that assesses the bottom-up activation of the reward system by conditioned stimuli. For more details on the paradigm see (Diekhof and Gruber 2010; Diekhof et al. 2012).

### fMRI Data Acquisition

fMRI was performed using a 3-Tesla Magnetom TIM Trio Siemens scanner (Siemens Healthcare, Erlangen, Germany) equipped with a standard eight-channel phased-array head coil. First, a T1-weighted anatomical data set with 1mm isotropic resolution was acquired. For fMRI, 31 axial slices were acquired in ascending direction (slice thickness=3mm; interslice gap=0.6mm) parallel to the anterior commissure–posterior commissure line, with a gradient-echo echo-planar imaging sequence (echo time 33 ms, flip angle 70°; field-of-view 192 mm, interscan repetition time 1900ms).

In 2 functional runs, 185 volumes each were acquired. Stimuli were viewed through goggles (Resonance Technology, Northridge, California, USA), and subjects responded via button presses on a fiber optic computer response device (Current Designs, Philadelphia, Pennsylvania, USA). To present the stimuli in the scanner, Presentation Software (Neurobehavioral Systems, Albany, California, USA) was used. Functional images were preprocessed and analyzed using SPM12 (https://www.fil.ion.ucl.ac.uk/spm/; SPM RRID:SCR_007037) with a general linear model. The study design was event-related. Only correctly answered trials were included in the analysis. Linear t-contrasts were defined to assess activation effects elicited by the conditioned reward stimuli.

### fMRI Phenotype Extraction

For the subsequent genetic association analyses, neuroimaging phenotypes were determined. Using SPM12, the individual mean beta estimates from the a priori regions of interest (VTA and NAc) were extracted. For each participant, beta extraction was performed using predefined MNI coordinates of both brain regions emerging from previous publications: MNI coordinates ±9 -21 -15 for the bilateral VTA and MNI coordinates ±12 12 -3 for the bilateral NAc (Diekhof et al. 2012; Jakob 2012; Trost et al. 2016). To account for interindividual functional neuroanatomical variation that may be insufficiently covered by spatial normalization processes in the standard preprocessing pipeline, a box with a 3x3x3mm dimension around the a priori MNI coordinates was applied within each subject to determine the individual maximum of reward-related brain activation.

### Genotyping, Variant Selection and Gene Set-Based Association Analysis

DNA was extracted from saliva, which was collected using Oragene DNA devices with standardized protocols from DNA Genotek (DNA Genotek, Ottawa, Ontario, Canada). Genotyping was performed using Illumina OmniExpress Genotyping BeadChips according to the protocols of the manufacturer (Illumina, Inc., San Diego, California, USA). Association testing was performed on imputed and quality-controlled genotype data (for details see (Treutlein et al. 2024)). Genotypes of the analyzed variants were in Hardy-Weinberg Equilibrium (HWE p>0.05) (Supplemental Table S1).

To reduce the number of required corrections due to multiple testing, gene set-based association analysis jointly tests genetic variants as a single gene set rather than as single markers. Genes coding for proteins functionally related to the LC3 conjugation system were selected from recent review articles (Zhou 2022; Levine and Kroemer 2019; Yamamoto et al. 2023). For analysis, missense genetic variants in the genes coding for these proteins were selected from the dbSNP database (Sherry et al. 1999)(https://www.ncbi.nlm.nih.gov/snp/; dbSNP database RRID:SCR_002338), using a global MAF setting of 0.001 to 1.0 (Table 1).

**Table 1.**
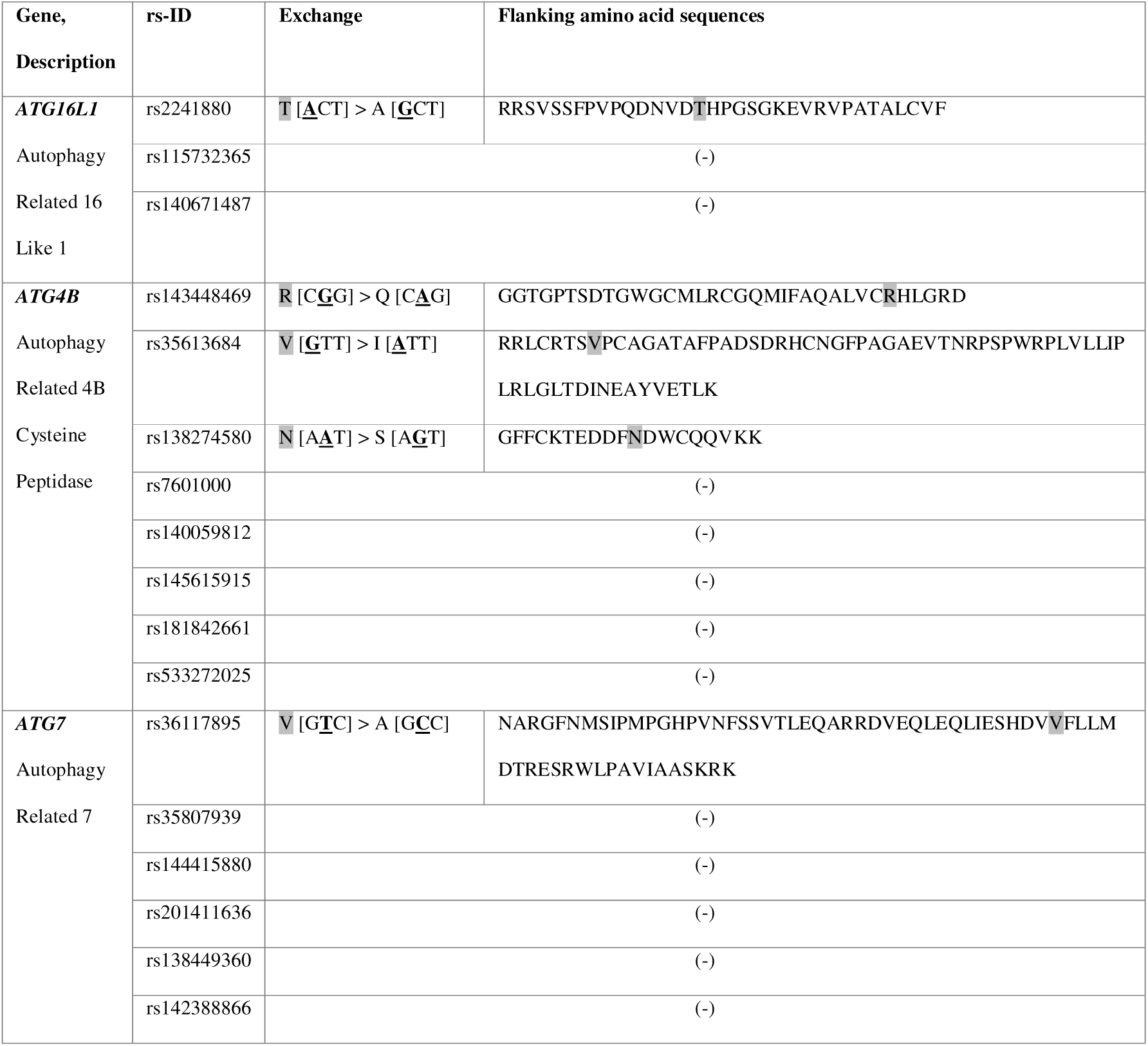
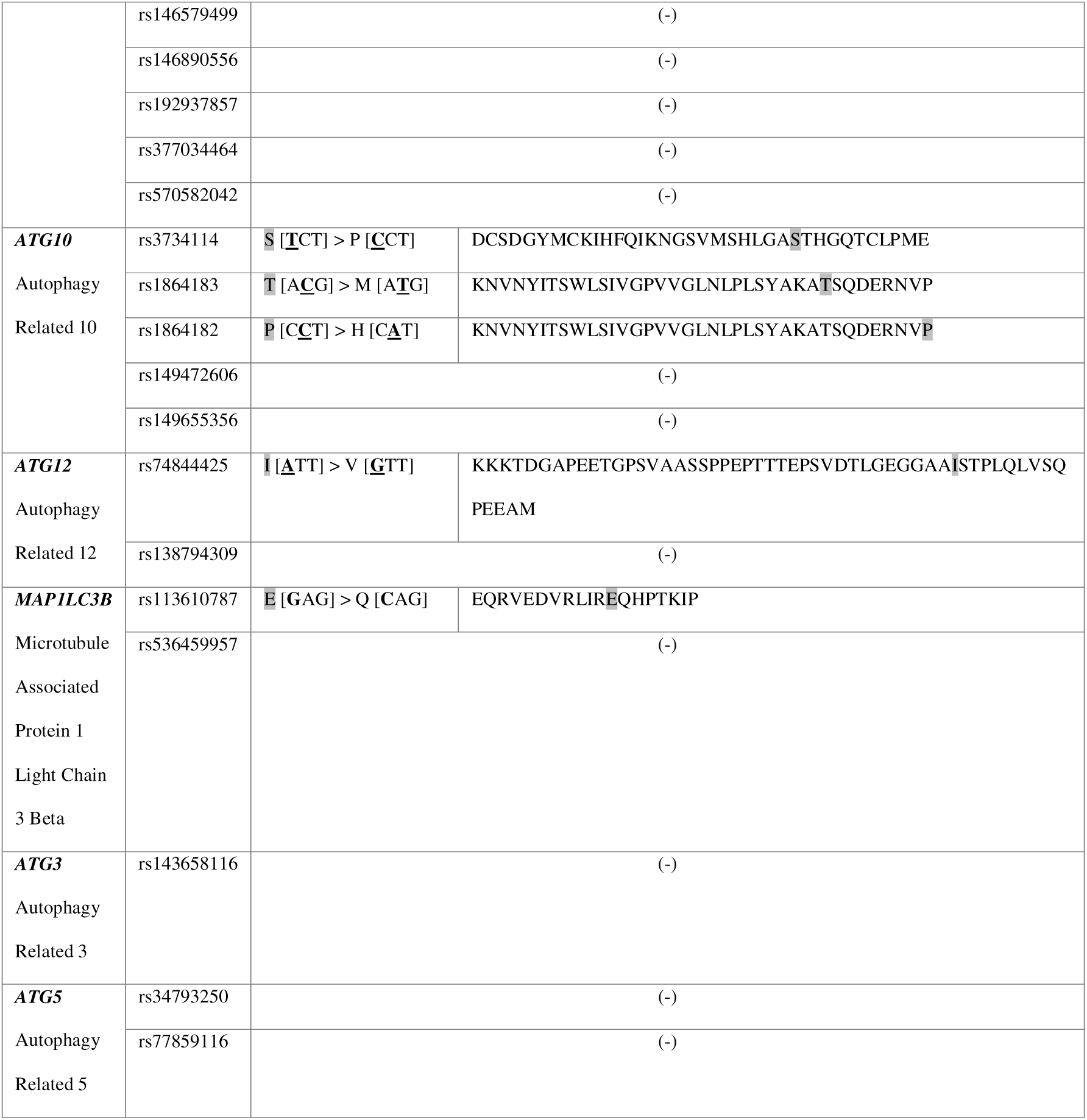
Selection of missense variants in genes related to the LC3 conjugation process. Gene: gene symbol; Description: gene name; rs-ID: rs-identifier of missense variants in dbSNP with global minor allele frequency of 0.001 to 1.0; Exchange: amino acid exchange and bold underlined the corresponding nucleotide exchange within the codon triplet, of missense variants from dbSNP which were present in the imputed GWAS dataset; (-): missense variant from dbSNP which was not present in the imputed dataset. Flanking amino acid sequences: flanking amino acid sequence of the exon (Haeussler et al. 2019) (UCSC Genome Browser database RRID:SCR_005780) with position of the amino acid exchange shaded grey.

In the gene set-based method used for the present study, the PLINK set-test, first an analysis of single variants in the original set is performed. From the statistics of the single variant analysis, a mean statistic is calculated. Then, the same analysis is performed with a certain number of simulated sets with a permuted phenotype status of the individuals. From these computations, an empirical P-value (’EMP1’) is retrieved by calculation of the number of times that the test statistic of the simulated sets exceeds the test statistic of the original set (Deelen et al. 2013). For the set-based association analysis, PLINK version v1.90b6.9 (Chang et al. 2015) (PLINK RRID:SCR_001757) was used with the adaptive procedure (flag ‘--assoc perm set-test’), without exclusion of any of the variants due to P-value threshold.

### Whole Brain Neuroimaging Analyses for MAP1LC3B rs113610787 (E25Q)

Neuroimaging analyses of the strongest associated variant in this study, *MAP1LC3B* rs113610787, were conducted to confirm the observed effects in subcortical structures, and for exploration of additional effects in other regions of the extended reward system. A full factorial design with the levels ‘major allele homozygotes’ (n=207) and ‘heterozygotes’ (n=7) was applied to assess genotypic effects on reward-related brain activation with a particular focus on the contrast ‘heterozygotes*>*major allele homozygotes’. Using SPM12, we performed second-level analyses based on first level single subjects contrast images related to brain activations elicited by the conditioned reward stimuli. Whole-brain genotype group associations were conducted with a search criterion of p<0.05, uncorrected.

### Detection of conservation at single amino-acid sites

Substitution rate variation across sites derive from differences in selective constraints (and thereby functional importance) on the sites, e.g., whether they participate in interactions between proteins, are involved in ligand binding, or are important for enzymatic activity. Rapidly evolving amino acid positions in a protein are variable, whereas slowly evolving positions are conserved (Mayrose et al. 2004; Pupko et al. 2002).

Orthologous protein sequences for MAP1LC3B were retrieved from DIOPT (https://www.flyrnai.org/cgi-bin/DRSC_orthologs.pl; DIOPT RRID:SCR_021963) (Hu et al. 2011) and multiple-sequence aligned using Clustal Omega (Sievers et al. 2011; Madeira et al. 2024) (https://www.ebi.ac.uk/jdispatcher/msa/clustalo; Clustal Omega RRID:SCR_001591), using default settings. Then, amino acid positions that contained gaps in the alignment were removed (Supplemental Figure S1). Conservation at single amino-acid sites were investigated with Rate4Site_v.2.01 (https://www.tau.ac.il/~itaymay/cp/rate4site.html; Rate4Site RRID:SCR_024222). Using a multiple sequence alignment and a given tree, Rate4Site computes the relative evolutionary rate for each individual site. The Newick format tree for Rate4Site analysis (without branch length; Supplemental Figure S2) corresponds to the placement of the model organism taxa in the tree of life (Ciccarelli et al. 2006). Rate4Site was run using the options -s seq.aln [MSA file] and -t tree.txt [tree file], all other options were default settings. The Rate4Site conservation score corresponds to the evolutionary rate of a site and informs about conservation properties as indicator of a sités functional constraint (Mayrose et al. 2004; Pupko et al. 2002; Sydykova and Wilke 2017).

## Results

### Gene Set-Based Association Analysis

Gene set-based analyses were used to investigate the joint contribution of missense variants in the LC3-PE conjugation process to the extent of activation of the reward system. For three regions of interest, we observed a significant joint effect of the variants in the gene set (EMP1_R-VTA_=0.008; EMP1_R-NAc_=0.009; EMP1_L-NAc_=0.039). Of these brain regions, R-VTA and R-NAc survived correction for four comparisons at p<0.05. Full gene set test results are shown in Table 2.

**Table 2.**
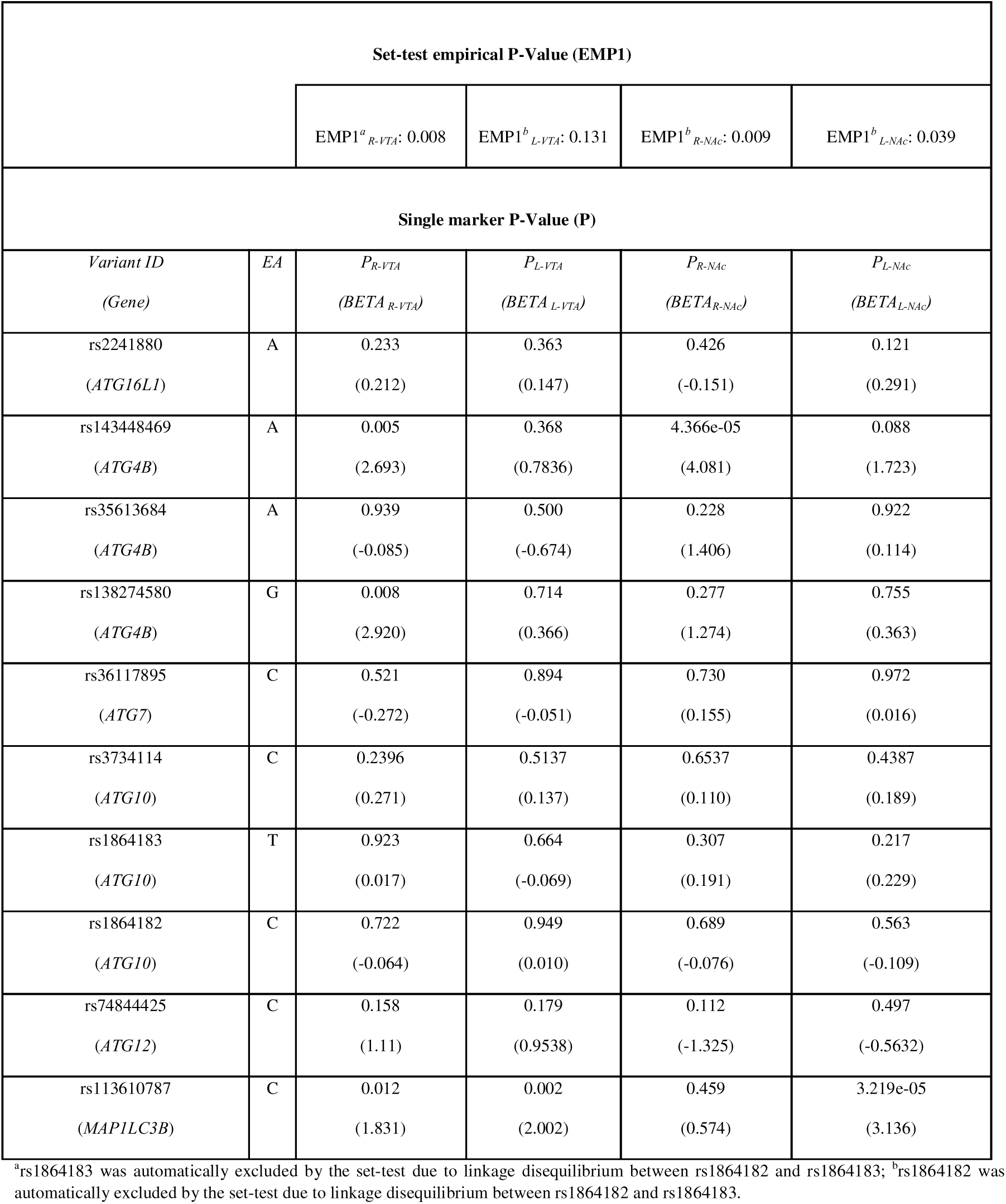
Set-based and single marker-based association results. For set-based analyses EMP1-values (set-test empirical P–values) are shown; for single marker analyses effect alleles (EA) and single marker P–values (P) with effect sizes (BETA) in parentheses are shown.

| Set-test empirical P-Value (EMP1) |  |  |  |  |  |
| --- | --- | --- | --- | --- | --- |
|  |  | EMP1<sup>a</sup><sub>R-VTA</sub>: 0.008 | EMP1<sup>b</sup><sub>L-VTA</sub>: 0.131 | EMP1<sup>b</sup><sub>R-NAc</sub>: 0.009 | EMP1<sup>b</sup><sub>L-NAc</sub>: 0.039 |
| Single marker P-Value (P) |  |  |  |  |  |
| Variant ID<br>(Gene) | EA | P<sub>R-VTA</sub><br>(BETA<sub>R-VTA</sub>) | P<sub>L-VTA</sub><br>(BETA<sub>L-VTA</sub>) | P<sub>R-NAc</sub><br>(BETA<sub>R-NAc</sub>) | P<sub>L-NAc</sub><br>(BETA<sub>L-NAc</sub>) |
| rs2241880<br>(ATG16L1) | A | 0.233<br>(0.212) | 0.363<br>(0.147) | 0.426<br>(-0.151) | 0.121<br>(0.291) |
| rs143448469<br>(ATG4B) | A | 0.005<br>(2.693) | 0.368<br>(0.7836) | 4.366e-05<br>(4.081) | 0.088<br>(1.723) |
| rs35613684<br>(ATG4B) | A | 0.939<br>(-0.085) | 0.500<br>(-0.674) | 0.228<br>(1.406) | 0.922<br>(0.114) |
| rs138274580<br>(ATG4B) | G | 0.008<br>(2.920) | 0.714<br>(0.366) | 0.277<br>(1.274) | 0.755<br>(0.363) |
| rs36117895<br>(ATG7) | C | 0.521<br>(-0.272) | 0.894<br>(-0.051) | 0.730<br>(0.155) | 0.972<br>(0.016) |
| rs3734114<br>(ATG10) | C | 0.2396<br>(0.271) | 0.5137<br>(0.137) | 0.6537<br>(0.110) | 0.4387<br>(0.189) |
| rs1864183<br>(ATG10) | T | 0.923<br>(0.017) | 0.664<br>(-0.069) | 0.307<br>(0.191) | 0.217<br>(0.229) |
| rs1864182<br>(ATG10) | C | 0.722<br>(-0.064) | 0.949<br>(0.010) | 0.689<br>(-0.076) | 0.563<br>(-0.109) |
| rs74844425<br>(ATG12) | C | 0.158<br>(1.11) | 0.179<br>(0.9538) | 0.112<br>(-1.325) | 0.497<br>(-0.5632) |
| rs113610787<br>(MAP1LC3B) | C | 0.012<br>(1.831) | 0.002<br>(2.002) | 0.459<br>(0.574) | 3.219e-05<br>(3.136) |

<sup>a</sup>rs1864183 was automatically excluded by the set-test due to linkage disequilibrium between rs1864182 and rs1864183; <sup>b</sup>rs1864182 was automatically excluded by the set-test due to linkage disequilibrium between rs1864182 and rs1864183.

### Single-Variant Association Analysis

In addition to computation of an empirical P-value of a gene set, PLINK 1.9 conducts association analyses of single markers contained in the set. In the two gene set tests that survived testing for multiple comparisons (R-VTA and R-NAc), three genetic variants reached significance at P<0.05: rs143448469 (P=0.005) and rs138274580 (P=0.008) in *ATG4B*, and rs113610787 (P=0.012) in *MAP1LC3B* were associated with the strength of activation in the R-VTA region, whereas rs143448469 (P=4.366e-05) in *ATG4B* influenced the strength of response in the R-NAc. Despite the set-test only showed a tendency to significance in the L-NAc, single marker rs113610787 in *MAP1LC3B* yielded a strong P=3.219e-05 for association with the response in this region. The same variant showed P=0.002 for the reactivity of the L-VTA, for which the set-test failed to reach the significance threshold at the 0.05 level.

Because rs143448469, rs138274580 and rs113610787 were involved in the association of two associated brain regions (R-VTA and R-NAc), their respective genes *ATG4B* and *MAP1LC3B* constitute the strongest findings of our analysis. Consistent across R-VTA and R-NAc brain regions, the risk allele of rs143448469 for increased activation was the minor A-allele, and the risk allele of rs113610787 for increased activation was the minor C-allele. Full single marker association test results for both brain regions and both samples are given in Table 2.

### Whole Brain Neuroimaging Analyses for MAP1LC3B rs113610787 (E25Q)

This analysis confirmed the already described effects of rs113610787 on reward-related activation in the left nucleus accumbens (NAc) (L-NAc: x=-9, y=12, z=0, T=3.11) (Figure 1a) and in the bilateral VTA (L-VTA: x=-3, y=-21, z=-12, T=3.43; R-VTA: x=3, y=-21, z=-12, T=3.71) (Figure 1b). In order to further explore the effects of rs113610787 in *MAP1LC3B* on other brain regions within the extended reward system (i.e., beside effects on our primary seed regions VTA and NAc), additional whole-brain group analyses were conducted comparing groups differing with respect to this genetic variant (contrast ‘heterozyotes>major allele homozygotes’). This analysis demonstrated that activation in further brain regions of the extended reward system, e.g., bilaterally in the in the intraparietal sulcus (IPS) (L-IPS: x=-36, y=-39, z=48, T=5.45; R–IPS: x=27, y=-45, z=45, T=3.34), inferior frontal sulcus (IFS) (L–IFS: x=-42, y=39, z=9, T=3.32; R-IFS: x=42, y=39, z=9, T=3.68), anteroventral prefrontal cortex (avPFC) (L-avPFC: x=-39, y=51, z=12, T=3.20; R–avPFC: x=27, y=54, z=6, T=4.41), anterior insula (L-AI: x=-36, y=24, z=-6, T=3.75; R-AI: x=39, y=15, z=0, T=4.24), and in the amygdala (L-AMY: x=-30, y=0, z=-27, T=3.56; R-AMY: x=30, y=0, z=-18, T=3.08). In the right frontomedian cortex (FMC) (R-FMC: x=6, y=18, z=60, T=4.13) and the left pregenual anterior cingulate cortex (L-pACC: x=-6, y=36, z=6, T=4.33) activation was also significantly increased in carriers of the rs113610787 minor allele.

**Figure 1.**
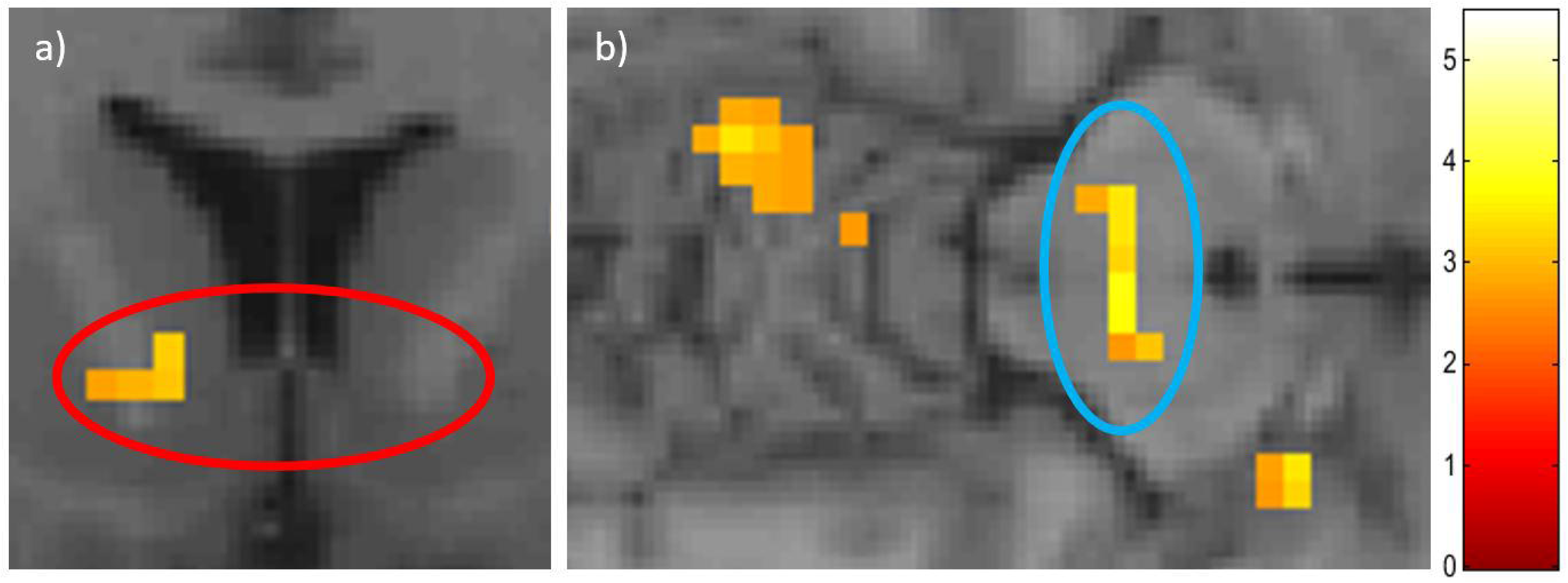
Effects of *MAP1LC3B* rs113610787 on activation of the a) ventral striatum at y=12, and of the b) ventral tegmental area at z=-12. Minor allele carriers show increased responses to conditioned reward stimuli as compared to major allele homozygotes. For illustration purposes, the T-map of these effects is shown at a threshold of p<0.005, uncorr

### Substitution Rate at the MAP1LC3B E25Q (rs113610787) site

*MAP1LC3B* rs113610787 (E25Q) codes for an exchange of the negatively charged amino acid glutamic acid (E) to the neutral amino acid glutamine (Q). The E25Q site is more variable (Rate4Site site rate: 0.910) compared to the Gly120 site (Rate4Site site rate: 0.047). At Gly120, which is invariant from human to yeast (Thukral et al. 2015)), the lipid is attached in the LC3 conjugation process, leading to a high functional constraint and a low position 120 site rate. The Rate4Site mean site rate across all MAP1LC3B sites was 0.498 ± 0.538 (min. 0.025, max. 2.998) (Supplemental Figure S1, Supplemental Figure S2).

## Discussion

The goal of our study was to test the hypothesis that allele-related functional variation in genes of the LC3 conjugation system is associated with the reactivity of subcortical key regions of the mesolimbic reward system in response to conditioned reward stimuli. The specific molecular process that was tested in the present study involves a chain of biochemical reactions that finally results in lipidation of autophagic protein LC3. Using a set-based test, we detected significant effects of relevant genes that contribute to LC3 conjugation in both analyzed brain regions. Particularly, *ATG4B* and *LC3B* emerged as factors that influenced the functional characteristics of the brain reward system.

ATG4B and LC3B of our gene set are components of the so called ATG4B-LC3 complex (Satoo et al. 2009), which was shown to be formed by the binding of the autophagy cystine protease ATG4B to LC3B with high affinity in vitro (Tang et al. 2022). ATG4B has a high catalytic efficiency to cleave the LC3B protein to generate LC3-I, which is conjugated with PE to form LC3-II, that attaches to phagophore membranes.

Previously published studies for emotional behavior in preclinical animal models, as well as in mood disorders in humans, showed that LC3 and ATG4 play key roles in these processes. Comprehensive support for correlation of LC3 and ATG4 with depression-like phenotypes (in several brain regions) derive from numerous preclinical stress models in rodents (reviewed in (Pierone et al. 2020)). In several depression-relevant animal models, signs of decreased autophagy has been reported (Gassen and Rein 2019); however decrease was not consistent across all reported depression-relevant animal models). For example in rodents, bacterial endotoxin lipopolysaccharide (LPS) treatment, an experimental model for behavioral alterations that resemble depression in humans, induced depression-like symptoms and changed expression of autophagic markers, among them protein LC3 in the central nervous system (Francois et al. 2014; Jiang et al. 2017).

Apart from preclinical studies in model organisms, LC3 also received support from a clinical study on the psychological status in major depressive disorder, as determined by the Symptom Distress Checklist (SCL-90-R), where LC3 gene expression in blood mononuclear cells was highly correlated with all nine subscales of the checklist. The high correlations observed in this study were interpreted by the authors as a biological connection between autophagy components, among them LC3 (correlation coefficients 0.522 – 0.862, P<0.001), and the clinical profile of depression (Alcocer-Gomez et al. 2017).

The relevance of the LC3 conjugation process, like the hypothesis of an involvement of the whole autophagic process in depression, derive to a substantial part from observations of antidepressant effects on autophagy (Gassen and Rein 2019). The majority of studies reported an induction of autophagy by antidepressants (Rein 2019), i.e. upon antidepressant treatment of cells, levels of autophagy markers rise (Gassen et al. 2015). Antidepressant drugs for which an effect on LC3 level have been described are, e.g., paroxetine, fluoxetine and amitryptiline. The clinical response to paroxetine in depressed patients could be predicted by the antidepressant response of autophagy markers, among them LC3B-II, in peripheral blood lymphocytes derived from patients, cultivated and treated ex vivo (Gassen et al. 2014). Another example of antidepressant effect on autophagy is the induction of autophagy by the antidepressive drugs fluoxetine and amitriptyline, via accumulation of sphingomyelin in lysosomes and Golgi membranes, and via ceramide in the endoplasmatic reticulum. Ceramide was shown to stimulate phosphatase 2A and thereby LC3B, among other autophagy proteins (Gulbins et al. 2018).

Due to the recognition that LC3 constitutes a druggable target with sufficient evidence and comprehensive structural information, this protein was suggested as candidate for future development of autophagosome modulators for various disease conditions (Costa and Ketteler 2015).

Behavioral response to rewards that is mediated by dopamine is a conserved trait as this neurotransmitter is already present in organisms that are evolutionary distant from humans. Its occurrence in roundworms, where dopamine modulates food seeking behavior, locomotor activity and reward-based learning, indicates that this neurotransmitter mediates these processes for more than one billion years (Nei et al. 2001; Costa and Schoenbaum 2022).

The MAP1LC3B E25Q site associated with reward system reactivity in the present study is neither conserved nor hypervariable. If the relaxed degree of conservation, as observed for the E25Q site, in further analyses will turn out to be a general feature of missense polymorphisms associated with reward system reactivity, a speculative explanatory model for this observation could be postulated: Strong conservation of reward-related behavioral processes could be advantageous for essential processes like feeding and sex, but could be disadvantageous in respect to other reward-related activities that require adaptive behavioral flexibility.

For example, investigation at the circuit-level in animals showed that the activity dynamics of the reward system encoded key features of social interaction (Gunaydin et al. 2014). The existence of a connection between social interaction and reward system activation is also evidenced by human fMRI studies (Kawamichi et al. 2016; Alkire et al. 2018; Krach et al. 2010). As social interaction depends on social contexts in groups (Yamaguchi et al. 2015), and can be expected to be subject to frequent change over evolutionary times, it may also relate to less conservation at the molecular level, potentially expressing as relaxed conservation of amino acid position site rate. Thus, from an evolutionary perspective, the E25Q site could be involved in adaptive modulation of the reward system.

To summarize the supporting evidences of our main finding, the ATG4B-LC3 complex was reported to play a key role in several stress-related conditions, possibly by impacting on oxidative stress and neuroinflammation, to influence synaptic function in multiple brain areas, potentially leading to depression-like phenotypes and psychiatric syndromes. By this mode of action, the ATG4B-LC3 complex may act on reward system reactivity in humans, the phenotype measured by us, which in turn may impact on progression to psychiatric disorders.

In sum, the present study investigated the molecular components of the LC3 conjugation process in the context of reward system reactivity in humans. Our findings indicate that particularly two genes that code for tightly interacting proteins in this process, *ATG4B* and *MAP1LC3B*, influence neural responses to conditioned reward stimuli. From the molecular systems perspective, the detected associations suggest that relevant genes related to the LC3 conjugation system of autophagy contribute to modulation of the mesolimbic dopaminergic reward system and in that way may also influence the vulnerability to neuropsychiatric disorders.

## Acknowledgements

We thank Prof. Dr. Elisabeth Binder and Monika Rex-Haffner, Max-Planck-Institute for Psychiatry, Munich for SNP genotyping and Maria Keil, Center for Translational Research in Systems Neuroscience and Psychiatry, Department of Psychiatry and Psychotherapy, Georg-August-University Göttingen, Göttingen, for support in recruitment and fMRI investigation of volunteers. We also thank Prof. Dr. Stephan Ripke and Swapnil Awasthi, Department of Psychiatry and Psychotherapy, Charité – Universitätsmedizin Berlin, Campus Mitte, Berlin, Germany, for genotype imputation and quality control of SNP genotype data. We also thank all subjects who participated in this study.

## Data availability statement

Data can be made available by the Principal Investigator (O.G.) upon reasonable request. The data are not publicly available due to privacy or ethical restrictions.

## Supplemental online material

Supplemental online material accompanies the manuscript.

