## Supplemental file for "Investigation of the Role of the LC3 Conjugation System in Autophagy for Human Reward System Reactivity"

**Supplemental Material**

**Supplemental Tables**

Table S1. Genotype-related information. Gene: gene symbol; Variant ID: rs-identifier; Chr: chromosome; BP: base position; Genotype counts: Major allele homozygotes / heterozygotes / minor allele homozygotes; MAF: minor allele frequency; HWE: Hardy-Weinberg equilibrium P-value.

| *Gene* | *Variant ID* | *Chr* | *BP* | *Genotype Counts* | *MAF*  *[Minor Allele]* | *HWE* |
| --- | --- | --- | --- | --- | --- | --- |
| *ATG16L1* | rs2241880 | 2 | 234183368 | 72/97/45 | 0.437 [A] | 0.267 |
| *ATG4B* | rs143448469 | 2 | 242593011 | 207/4/0 | 0.009 [A] | 1.0 |
| *ATG4B* | rs35613684 | 2 | 242606080 | 210/3/0 | 0.007 [A] | 1.0 |
| *ATG4B* | rs138274580 | 2 | 242610172 | 211/3/0 | 0.007 [G] | 1.0 |
| *ATG7* | rs36117895 | 3 | 11400019 | 195/18/1 | 0.047 [C] | 0.372 |
| *ATG10* | rs3734114 | 5 | 81354389 | 135/71/8 | 0.203 [C] | 0.835 |
| *ATG10* | rs1864183 | 5 | 81549216 | 60/98/56 | 0.491 [T] | 0.220 |
| *ATG10* | rs1864182 | 5 | 81549240 | 69/99/46 | 0.447 [C] | 0.337 |
| *ATG12* | rs74844425 | 5 | 115177207 | 208/6/0 | 0.014 [C] | 1.0 |
| *MAP1LC3B* | rs113610787 | 16 | 87432453 | 207/7/0 | 0.016 [C] | 1.0 |

**Supplemental Figures**

Figure S1. Multiple sequence alignment of MAP1LC3B orthologs of eukaryotic model organisms, for computation of Rate4Site amino acid position site rates. MAP1LC3B orthologous protein sequences from DIOPT (Hu et al 2011) were used for multiple sequence alignment in Clustal Omega (Sievers et al. 2011; Madeira et al. 2024). The amino acid position of MAP1LC3B E25Q (rs113610787) is underlined and the position of the Gly120 site is shaded grey; Positions that contain gaps are highlighted black and were removed prior to Rate4Site analysis; Human: *Homo sapiens*; Mouse: *Mus musculus*; Rat: *Rattus norvegicus*; Frog (Western clawed frog): *Xenopus tropicalis*; Fish (Zebrafish): *Danio rerio*; Fly: *Drosophila melanogaster*; Mosquito: *Anopheles gambiae*; Yeast: *Saccaromyces cerevisiae*; Fission yeast: *Schizosaccaromyces pombe*; ThCr (Thale cress): *Arabidopsis thaliana*. #: position numbering according to Rate4Site output in Figure S2.

```
                     00000000000000000000000000000000000000000000000000
                     00000000011111111112222222222333333333344444444445
                     12345678901234567890123456789012345678901234567890 Human_MAP1LC3B       MPSEKTFKQRRTFEQRVEDVRLIREQHPTKIP------------------
Mouse_Map1lc3b       MPSEKTFKQRRSFEQRVEDVRLIREQHPTKIPVGSPAARGDPNPIGAPRS
Rat_Map1lc3b         MPSEKTFKQRRSFEQRVEDVRLIREQHPTKIP------------------
Frog_map1lc3a        MPSERPFKHRRTFAERCAEVRQIREQHPNKIP------------------
Fish_map1lc3b        MPSEKTFKQRRTFEQRVEDVRLIREQHPNKIP------------------
Worm_lgg_2           -----SFKERRPFHERQKDVEEIRSQQPNKVP------------------
Fly_Atg8b            ------YKKDHSFDKRRNEGDKIRRKYPDRVP------------------
Mosquito_AgaP_AGAP00 ------YKEEHPFEKRKAEGDKIRRKYPDRVP------------------
Yeast_ATG8           ---KSTFKSEYPFEKRKAESERIADRFKNRIP------------------
Fission_Yeast_atg8   ------FKDDFSFEKRKTESQRIREKYPDRIP------------------
ThCr_APG8H           ----KSFKEQYTLDERLAESREIIAKYPTRIP------------------
ThCr_ATG8H           ----KSFKDQFSSDERLKESNNIIAKYPDRIP------------------
#                                    1111111111222222
#                          01234567890123456789012345


                     00000000000000000000000000000000000000000000000001
                     55555555556666666667777777777888888888899999999990
                     12345678901234567890123456789012345678901234567890 Human_MAP1LC3B       -----------------VIIERYKGEKQLPVLDKTKFLVPDHVNMSELIK
Mouse_Map1lc3b       PGKPQAPDLSRGKRGRRVIIERYKGEKQLPVLDKTKFLVPDHVNMSELIK
Rat_Map1lc3b         -----------------VIIERYKGEKQLPVLDKTKFLVPDHVNMSELIK
Frog_map1lc3a        -----------------VIIERYKGEKQLPVLDKTKFLVPDHVNMSELVK
Fish_map1lc3b        -----------------VIIERYKGEKQLPILDKTKFLVPDHVNMSELIK
Worm_lgg_2           -----------------VIIERFDGERSLPLMDRCKFLVPEHITVAELMS
Fly_Atg8b            -----------------VIVEKA-PKTRYAELDKKKYLVPADLTVGQFYF
Mosquito_AgaP_AGAP00 -----------------VIVEKA-PKARIDDLDKKKYLVPSDLTVGQFYF
Yeast_ATG8           -----------------VICEKA-EKSDIPEIDKRKYLVPADLTVGQFVY
Fission_Yeast_atg8   -----------------VICEKV-DKSDIAAIDKKKYLVPSDLTVGQFVY
ThCr_APG8H           -----------------VIAEKY-CKTDLPAIEKKKFLVPRDMSVGQFIY
ThCr_ATG8H           -----------------VIIEKYS-NADLPDMEKNKYLVPRDMTVGHFIH
#                                     222233  3333333344444444445555555
#                                     678901  2345678901234567890123456


                     11111111111111111111111111111111111111111111111111
                     00000000011111111112222222222333333333344444444445
                     12345678901234567890123456789012345678901234567890 Human_MAP1LC3B       IIRRRLQLNANQAFFLLVNGHSMVSVSTPISEVYESEKDEDGFLYMVYAS
Mouse_Map1lc3b       IIRRRLQLNANQAFFLLVNGHSMVSVSTPISEVYESERDEDGFLYMVYAS
Rat_Map1lc3b         IIRRRLQLNANQAFFLLVNGHSMVSVSTPISEVYESERDEDGFLYMVYAS
Frog_map1lc3a        IIRRRLQLNPTQAFFLLVNQHSMVSVSTPILDIYEQEKDEDGFLYMVYAS
Fish_map1lc3b        IIRRRLQLNSNQAFFLLVNGHSMVSVSTAISEVYERERDEDGFLYMVYAS
Worm_lgg_2           IVRRRLQLHPQQAFFLLVNERSMVSNSMSMSNLYSQERDPDGFVYMVYTS
Fly_Atg8b            LIRKRINLRPDDALFFFVN-NVIPPTSATMGALYQEHFDKDYFLYISYTD
Mosquito_AgaP_AGAP00 LIRKRIHLRPEDALFFFVN-NVIPPTSATMGSLYHEHHEEDYFLYIAYSD
Yeast_ATG8           VIRKRIMLPPEKAIFIFVN-DTLPPTAALMSAIYQEHKDKDGFLYVTYSG
Fission_Yeast_atg8   VIRKRIKLSPEKAIFIFID-EILPPTAALMSTIYEEHKSEDGFLYITYSG
ThCr_APG8H           ILSARLHLSPGKALFVFVN-NTLPQTAALMDSVYESYKDDDGFVYMCYSS
ThCr_ATG8H           MLSKRMQLDPSKALFVFVH-NTLPQTASRMDSLYNTFKEEDGFLYMCYST
#                                                                111111
#                    5556666666666777777 777788888888889999999999000000
#                    7890123456789012345 678901234567890123456789012345


                     1111111111
                     5555555556
                     1234567890
Human_MAP1LC3B       QETFGMKLSV
Mouse_Map1lc3b       QETFGTAMAV
Rat_Map1lc3b         QETFGTALAV
Frog_map1lc3a        QETFG-----
Fish_map1lc3b        QETFGFQ---
Worm_lgg_2           QPAFG-----
Fly_Atg8b            ENVYG-----
Mosquito_AgaP_AGAP00 ENVYG-----
Yeast_ATG8           ENTFG-----
Fission_Yeast_atg8   ENTFG-----
ThCr_APG8H           EKTFG-----
ThCr_ATG8H           EKTFG-----
                     11111
                     00001
                     67890
```

Figure S2. Amino acid position site rates from Rate4Site analysis of MAP1LC3B orthologs in eukaryotic model organisms. Parameters of the analysis were determined by Rate4Site (Pupko et al. 2002, Mayrose et al. 2004). Newick format tree: ((((((((Mouse,Rat),Human),Frog),Fish),(Mosquito,Drosophila)),Worm),(Yeast,Fission_Yeast)),(ThCr1,ThCr2)); The amino acid position of MAP1LC3B E25Q (rs113610787) is underlined and the position of the Gly120 site is shaded grey, relating to the multiple sequence alignment in Figure S1.

THE PARAMETERS

rate inference method is: empirical Bayesian estimate using a Gamma prior distribution with: 16 discrete categories

probablistic_model is: JTT

branch lengths optimization is ML using a gamma model

Computing the rates...

rate of pos: 0 = 0.16911

rate of pos: 1 = 0.035177

rate of pos: 2 = 1.4308

rate of pos: 3 = 0.794101

rate of pos: 4 = 0.699307

rate of pos: 5 = 1.35869

rate of pos: 6 = 0.31569

rate of pos: 7 = 0.747061

rate of pos: 8 = 0.626756

rate of pos: 9 = 0.0306902

rate of pos: 10 = 1.93186

rate of pos: 11 = 2.99801

rate of pos: 12 = 0.300265

rate of pos: 13 = 0.279744

rate of pos: 14 = 1.2295

rate of pos: 15 = 1.90731

rate of pos: 16 = 0.0254724

rate of pos: 17 = 0.283852

rate of pos: 18 = 0.910315

rate of pos: 19 = 0.361212

rate of pos: 20 = 0.485961

rate of pos: 21 = 0.157297

rate of pos: 22 = 0.615475

rate of pos: 23 = 0.202146

rate of pos: 24 = 0.162885

rate of pos: 25 = 0.0419358

rate of pos: 26 = 0.0262708

rate of pos: 27 = 0.0254724

rate of pos: 28 = 0.585336

rate of pos: 29 = 0.0340236

rate of pos: 30 = 0.207447

rate of pos: 31 = 0.970431

rate of pos: 32 = 0.402497

rate of pos: 33 = 0.896806

rate of pos: 34 = 0.472277

rate of pos: 35 = 0.528076

rate of pos: 36 = 0.485306

rate of pos: 37 = 2.75045

rate of pos: 38 = 0.418909

rate of pos: 39 = 0.121862

rate of pos: 40 = 0.129113

rate of pos: 41 = 1.00708

rate of pos: 42 = 0.035177

rate of pos: 43 = 0.453112

rate of pos: 44 = 0.0471876

rate of pos: 45 = 0.0262708

rate of pos: 46 = 0.0419358

rate of pos: 47 = 1.11055

rate of pos: 48 = 0.245328

rate of pos: 49 = 0.45645

rate of pos: 50 = 0.181078

rate of pos: 51 = 0.106246

rate of pos: 52 = 0.335016

rate of pos: 53 = 0.317246

rate of pos: 54 = 0.311355

rate of pos: 55 = 0.601914

rate of pos: 56 = 0.935699

rate of pos: 57 = 0.344701

rate of pos: 58 = 0.228216

rate of pos: 59 = 0.110281

rate of pos: 60 = 0.341469

rate of pos: 61 = 0.0306902

rate of pos: 62 = 0.466707

rate of pos: 63 = 0.729554

rate of pos: 64 = 0.0471876

rate of pos: 65 = 0.85972

rate of pos: 66 = 0.308493

rate of pos: 67 = 1.57904

rate of pos: 68 = 0.28366

rate of pos: 69 = 0.0254841

rate of pos: 70 = 0.454929

rate of pos: 71 = 0.0481076

rate of pos: 72 = 0.378845

rate of pos: 73 = 0.311355

rate of pos: 74 = 0.102155

rate of pos: 75 = 0.158143

rate of pos: 76 = 0.60601

rate of pos: 77 = 0.319788

rate of pos: 78 = 0.222768

rate of pos: 79 = 0.419926

rate of pos: 80 = 0.334624

rate of pos: 81 = 0.211034

rate of pos: 82 = 0.0861495

rate of pos: 83 = 0.380856

rate of pos: 84 = 1.08171

rate of pos: 85 = 0.0918624

rate of pos: 86 = 0.400101

rate of pos: 87 = 1.37596

rate of pos: 88 = 0.754671

rate of pos: 89 = 0.0504627

rate of pos: 90 = 1.3763

rate of pos: 91 = 1.58188

rate of pos: 92 = 0.70732

rate of pos: 93 = 0.727485

rate of pos: 94 = 0.432729

rate of pos: 95 = 1.19028

rate of pos: 96 = 0.0306638

rate of pos: 97 = 0.26608

rate of pos: 98 = 0.0481076

rate of pos: 99 = 0.400575

rate of pos: 100 = 0.0504627

rate of pos: 101 = 0.26638

rate of pos: 102 = 0.639923

rate of pos: 103 = 0.0504627

rate of pos: 104 = 0.287183

rate of pos: 105 = 0.400104

rate of pos: 106 = 0.215416

rate of pos: 107 = 0.655852

rate of pos: 108 = 0.196185

rate of pos: 109 = 0.16911

rate of pos: 110 = 0.0472721

number of sites: 111
